# Call for participation: Collaborative benchmarking of functional-structural root architecture models. The case of root water uptake

**DOI:** 10.1101/808972

**Authors:** Andrea Schnepf, Christopher K. Black, Valentin Couvreur, Benjamin M. Delory, Claude Doussan, Axelle Koch, Timo Koch, Mathieu Javaux, Magdalena Landl, Daniel Leitner, Guillaume Lobet, Trung Hieu Mai, Félicien Meunier, Lukas Petrich, Johannes A. Postma, Eckart Priesack, Volker Schmidt, Jan Vanderborght, Harry Vereecken, Matthias Weber

**Affiliations:** Institut für Bio- und Geowissenschaften: Agrosphäre (IBG-3), Forschungszentrum Jülich GmbH, Wilhelm-Johnen-Str., D-52425 Jülich, Germany; Department of Plant Science, The Pennsylvania State University, 102 Tyson Building, University Park PA 16802, USA; Earth and Life Institute, Agronomy, Université catholique de Louvain, Louvain-la-Neuve, Belgium; Institute of Ecology, Leuphana University Lüneburg, Universitätsallee 1, 21335 Lüneburg, Germany; UMR 1114 EMMAH, INRA/UAPV, 84914, Avignon cedex 9, France; Earth and Life Institute, Environmental Sciences, Université catholique de Louvain, Louvain-la-Neuve, Belgium; Department of Hydromechanics and Modelling of Hydrosystems, University of Stuttgart, Pfaffenwaldring 61, 70569 Stuttgart, Germany; Simulationswerkstatt, Ortmayrstrasse 20, A-4060 Leonding, Austria; CAVElab - Computational and Applied Vegetation Ecology, Ghent University, Ghent, Belgium; Department of Earth and Environment, Boston University, Boston, USA; Institute of Stochastics, Ulm University, Helmholtzstr. 18, D-89069 Ulm, Germany; Institut für Bio- und Geowissenschaften: Plant Sciences (IBG-2), Forschungszentrum Jülich GmbH, Wilhelm-Johnen-Str., D-52425 Jülich, Germany; Institute of Soil Ecology, Helmholtz Zentrum München, Ingolstädter Landstr. 1, 85764 Neuherberg, Germany; International Soil Modelling Consortium ISMC, Jülich, Germany

## Abstract

Three-dimensional models of root growth, architecture and function are becoming important tools that aid the design of agricultural management schemes and the selection of beneficial root traits. However, while benchmarking is common in many disciplines that use numerical models such as natural and engineering sciences, functional-structural root architecture models have never been systematically compared. The following reasons might induce disagreement between the simulation results of different models: different representation of root growth, sink term of root water and solute uptake and representation of the rhizosphere. Presently, the extent of discrepancies is unknown, and a framework for quantitatively comparing functional-structural root architecture models is required. We propose, in a first step, to define benchmarking scenarios that test individual components of complex models: root architecture, water flow in soil and water flow in roots. While the latter two will focus mainly on comparing numerical aspects, the root architectural models have to be compared at a conceptual level as they generally differ in process representation. Therefore defining common inputs that allow recreating reference root systems in all models will be a key challenge. In a second step, benchmarking scenarios for the coupled problems are defined. We expect that the results of step 1 will enable us to better interpret differences found in step 2. This benchmarking will result in a better understanding of the different models and contribute towards improving them. Improved models will allow us to simulate various scenarios with greater confidence and avoid bugs, numerical errors or conceptual misunderstandings. This work will set a standard for future model development.

## 1 Introduction

A growing number of different modelling techniques and software libraries are now available to build functional-structural root architecture models. Different available models of root architecture and functions have been discussed and qualitatively compared in Dunbabin et al. (2013). The available models differ in the way they represent different processes such as root growth, water flow, solute transport are captured and translated into mathematical equations (process-level differences); in how they solve mathematical problems by their choice of analytical or numerical approach, numerical scheme, programming technique (solution-level differences); and in how they couple the different processes to the full model (coupling-level differences). However, the extent of discrepancies is currently unknown. Thus, a framework for quantitatively comparing functional-structural root architecture models is required. In addition to the explanatory or predictive power of a model, it is also important to understand the performance of these models, e.g. in terms of accuracy or computational cost. The most commonly used type of functional-structural root architecture models represent the structure of the root system as a 1-dimensional branched network of discrete segments which is geometrically embedded in a 3-dimensional soil domain (Koch et al., 2018b). The root architecture may either be known from measurements, such as 2D or 3D images, or from root architectural models. Suitable models are then used to simulate the “functions”, such as carbon flow and use in root systems (Bidel et al., 2000, e.g.), rhizodeposition (Nygren and Perttunen, 2010), competition between species (Dun), plant anchorage (Dupuy et al., 2007), water and nutrient uptake (Dunbabin et al., 2006; Javaux et al., 2008). Exchange between soil and root is typically modelled via source/sink terms. From the point of view of the soil domain, roots are often considered as line sources, i.e. it is assumed that their diameter is small compared to the relevant spatial scale of the soil. The advantage of this approach is that it does allow to consider root system architecture (position of each segment in time and 3D space) explicitly while being computationally less expensive than an explicit representation of root volumes in the soil domain. By direct comparison with explicit 3D simulations, Daly et al. (2017) showed that the error made by neglecting root volumes physically present in the soil domain is negligibly small in case of root water uptake. Thus, models of this type are sufficiently accurate and computationally cheaper than explicit 3D. The challenge is now to develop a commonly accepted framework for benchmarking functional-structural root architecture models. This includes defining a set of benchmark problems to test model accuracy and performance. We propose that models should be evaluated against two different kinds of references: First, we will develop simple benchmark scenarios, if possible with analytical solutions, that serve as a reference for model verification. Secondly, we define data sets that can be used as references for the evaluation of more complex models without analytical solution. These data sets should as good as possible describe the system we want to model and contain as little uncertainty as possible (Luo et al., 2012). This benchmark activity focuses on two processes, root architecture development and root water uptake. We propose this benchmarking framework to be used by the community of modellers and other participants to compare their model outputs against those of the reference solutions of benchmarks defined in this paper. The use of this framework thus aims to be a collaborative effort. We will refer to any numerical model that implemented some or all of the benchmark problems as “participating model” or “simulator”.

## 2 Benchmark problems for models of root architecture and function

In order to benchmark models of root architecture and function, we propose a multi-step approach with growing level of complexity.The individual benchmarks refer as much as possible to published work, however, we streamlined the different problems and made the notation consistent throughout this paper. A list of symbols is provided in Table 1. The intrinsic nature of functional-structural root architecture models involves multiple coupled domains and processes. A single process in a single domain (e.g. water flow in soil) is referred to as “module” here. The first set of benchmarks (M1-M3) is about individual modules (M) only, i.e. they either deal with only root growth, water flow in soil or water flow inside roots. The scenarios are simple, possibly have analytical solutions, and the goal is to build trust in the accuracy of the individual participating models and to help interpret potentially diverging results of the coupled benchmark problems. Benchmark problems M1 are about root architecture development. It is known that the representation of growth processes can be very different between different simulators. Thus, the goal is to calibrate each simulator individually to given root image data (reference data). M2 is about modelling water flow in soil. Here all participating models solve the same equation, namely the Richards equation, and differences may occur due to differences in numerical implementation. M3 deals with water flow inside the root system for static soil water conditions. As for M2, differences between models are expected to be mainly due to the numerical implementation of this well defined process. The second set of benchmarks (C1 and C2) is about coupled root-soil models. Benchmark problems C1 consider a static (non-growing) root system and focus on comparison of numerical representation of agreed-upon equations and process representations as well as on the coupling approach to compute the sink term for root water uptake. For this benchmark, we provide a reference solution that is based on a computational mesh that was generated with consideration of the physical presence of the roots in the soil domain. Thus, root water uptake was simulated not by a sink term but as a boundary condition at the root surface in soil. Our approach is similar to Daly et al. (2017) but in addition couples the soil domain to the root domain so that pressure gradients along the roots are simulated. Benchmark problem C2 compares the water uptake of fully coupled models with growing root systems.

**Table 1:** List of notations.

| Symbol | Units | Description |
| --- | --- | --- |
| $d$ | cm | depth |
| $D_w$ | $\text{cm}^2 \text{d}^{-1}$ | water diffusivity |
| $\mathbf{e}_3$ | (0,0,1) | standard unit vector |
| $J$ | $\text{cm}^3 \text{cm}^{-2} \text{d}^{-1}$ | water flux per unit soil surface area |
| $k_r$ | $\text{cm}^3 \text{cm}^{-2} \text{cm}^{-1} \text{d}^{-1}$ | root radial conductivity (defined as volume of water per unit root surface area, pressure head gradient and time) |
| $k_x$ | $\text{cm}^4 \text{cm}^{-1} \text{d}^{-1}$ | specific root axial conductance |
| $K(\theta)$ | $\text{cm}^3 \text{cm}^{-2} \text{d}^{-1}$ | soil hydraulic conductivity |
| $K_{sat}$ | $\text{cm}^3 \text{cm}^{-2} \text{d}^{-1}$ | saturated soil hydraulic conductivity |
| $l$ | cm | length |
| $n$ | - | van Genuchten shape parameter |
| $q$ | $\text{cm}^3 \text{cm}^{-2} \text{d}^{-1}$ | water flux per unit root surface area |
| $Q$ | $\text{cm}^3 \text{d}^{-1}$ | volumetric water flow rate |
| $\overline{Q}$ | $\text{cm}^3 \text{d}^{-1}$ | daily average volumetric water flow rate |
| $Q_r$ | $\text{cm}^3 \text{d}^{-1}$ | radial root water flow rate |
| $Q_x$ | $\text{cm}^3 \text{d}^{-1}$ | axial root water flow rate |
| $r_{root}$ | cm | root radius |
| $S_w$ | $\text{cm d}^{-0.5}$ | sorptivity (infiltration) or desorptivity (evaporation) |
| $t$ | d | time |
| $\mathbf{v}$ | $(v_1, v_2, v_3)$ | normalised direction of the xylem, pointing towards the root tip |
| $w$ | cm | width |
| $x, y, z$ | | spatial coordinates, z-axis pointing upward, soil surface is at $z=0$ |
| $Y$ | - | cumulative root fraction from surface to depth $d$ |
| $\alpha$ | $\text{cm}^{-1}$ | van Genuchten shape parameter |
| $\beta$ | - | root distribution index |
| $\eta$ | cm | position of the infiltration front (eqn. (4)) |
| $\lambda$ | - | van Genuchten-Mualem parameter |
| $\Lambda$ | - | root domain (network of root center-lines) |
| $\Omega$ | - | soil domain |
| $\Phi$ | $\text{cm}^2 \text{d}^{-1}$ | matric flux potential |
| $\theta$ | $\text{cm}^3 \text{cm}^{-3}$ | volumetric water content |
| $\theta_a$ | $\text{cm}^3 \text{cm}^{-3}$ | reference water content |
| $\theta_{res}$ | $\text{cm}^3 \text{cm}^{-3}$ | residual water content |
| $\theta_{sat}$ | $\text{cm}^3 \text{cm}^{-3}$ | saturated water content |
| $\psi$ | cm | water pressure head, described as potential energy per unit weight of water (i.e. units are cm of water column), given as relative to air pressure of 1020 cm and excluding the gravitational potential |
| $\zeta$ | | local coordinate along root axis |
| **sub indices** |  |  |
| collar | root collar (upper boundary of root system domain) |  |
| i | initial |  |
| pot | potential |  |
| r | radial |  |
| res | residual |  |
| s | soil |  |
| sat | saturation |  |
| seg | root segment |  |
| sim | simulation |  |
| sur | soil surface |  |
| tip(s) | root tip(s) (boundaries of root system domain) |  |
| top | top, position of the soil surface |  |
| out | outer radius of soil cylinder around a single root |  |
| x | xylem |  |

Each benchmark problem is described in a Jupyter Notebook that is publicly available on a github repository. We will provide codes for automatic analyses and comparison of different model results with the reference solutions or reference data. This makes the analysis transparent and easily modifiable and facilitates including even future participating models’ outputs at any later time.

### Levels of contribution

Any group using or developing functional-structural root architecture models is invited to participate in this collaborative model comparison. Not every model might be suited for all of the provided benchmark problems. Thus, every participant may decide in which individual benchmark problem they would like to participate. However, to reach a certain level of complexity, the “module” benchmarks should be simulated first before the “coupled” benchmarks. Table 2 gives an overview of the key features of these problems and their implementations. One important aim of this activity is a joint publication that shows and discusses the results of the different participating models in comparison to the reference solutions and reference data provided as well as to gain an overview of the extent of deviations between the different simulators.

**Table 2:** Description of benchmark scenarios to be implemented in 3D functional-structural root architecture models.^1^.

|  | Benchmark problem | Domain | Initial conditions | Boundary conditions | Evaluation | Remarks |
| --- | --- | --- | --- | --- | --- | --- |
| RSA | M1.1: RSA calibration | $t_{sim}=11$ (8) for lupine (maize) | seed position (0,0,-3) | n.a. | Comparison against the measured root systems provided - traits and persistent homology (PH) | Model parameters are determined from calibration against traced images provided in the github repository in RSML format in the folder in **M1.1 RSA calibration/M1.1 Reference data**; 100 realisations for each model setup |
| | M1.2: RSA simulation | $t_{sim}=60$ | seed position (0,0,-3) | n.a. | No reference solution, comparison amongst models - traits, PH, RLD | RSA model parameters from M1.1; 10 realisations for each model setup |
| Soil | M2.1: Infiltration | $1 \times w \times d = 10 \times 10 \times 200$, $t_{sim}=1$ | $\psi_{s,i} = -400$ | at $z = 0 \begin{cases} J_s = 100 \text{ if } \psi_s < 0 \\ \psi_s = 0 \text{ else} \end{cases}$,<br>$\frac{\partial \psi_s}{\partial z}\vert _{z=200} = 1$, no-flux at the sides | Analytical solution, eqn. (3) | sand, loam, clay (Table 3) |
| | M2.2: Evaporation | $1 \times w \times d = 10 \times 10 \times 100$, $t_{sim}=10$ | $\psi_{s,i} = -40$ for sand and $-200$ for all other scenarios | at $z = 0 \begin{cases} J_s = J_{s,ref} \text{ if } \psi_s > -10,000 \\ \psi_s = -10,000 \text{ else} \end{cases}$,<br>no-flux at all other boundaries | Analytical solution, eqn. (4) | scenario 1: sand, $J_{s,ref} = -0.1$,<br>scenario 2: loam, $J_{s,ref} = -0.1$,<br>scenario 3: loam, $J_{s,ref} = -0.3$,<br>scenario 4: clay, $J_{s,ref} = -0.3$ |
| Xylem | M3.1: Single root | 1 vertical root, $L=50$ | n.a. | $\psi_x\vert _{collar} = -1000$, $Q_r\vert _{tip} = 0$ | Analytical solution, Eq. (11) | $k_x=0.0432$, $k_r=1.73 \times 10^{-4}$, $\psi_s = -200$ |
| | M3.2: Root system | 14-day old root system | n.a. | $\psi_x\vert _{collar} = -500$, $Q_r\vert _{tips} = 0$ | Hybrid analytical solution (Meunier et al., 2017) | root hydraulic properties in scenario (a): Table 4, (b): Fig. 7, $\psi_s = -200$, static RSA given in the **root_grid** folder of this benchmark |
| Coupled 1 | C1.1: Single RWU | 1D radially symmetric, $r_{root} = 0.02$, $r_{out} = 0.6$, $t_{sim}=20$ | $\psi_{s,i} = -100$ | at $r = r_{root} \begin{cases} q_r = q_{root}, \text{ if } \psi_s > -15,000 \\ \psi_s = -15,000 \text{ else} \end{cases}$,<br>$q_r\vert _{r=r_{out}} = 0$ | Analytical solution, Eqs. (16) and (17) | sand, loam, clay (Table 3), scenarios 1-3: $q_{root}=0.1$, scenarios 4-6: $q_{root}=0.05$ |
| | C1.2: RWU, static RSA | static 8-day old root system, soil: $1 \times w \times d = 8 \times 8 \times 15$, $t_{sim}=3$ | $\psi_{s,i} = -659.8 - z$ | $Q_r\vert _{collar} = \begin{cases} Q_r = 6.4 \text{ if } \psi_s > -15,290 \\ \psi_s = -15,290 \text{ else} \end{cases}$,<br>$Q_r\vert _{tips} = 0$, no-flux at all soil faces | Reference solution: explicit 3D simulation | loam (Table 3), static RSA given in the **root_grid** folder of this benchmark, root hydraulic properties in scenario (a): Table 4, (b): Fig. 7 |
| Coupled 2 | C2.1: RWU, dynamic RSA | growing root system, soil: $1 \times w \times d = 25 \times 25 \times 100$, $t_{sim}=60$ | $\psi_{s,i} = -200$ | $Q_r\vert _{collar} = \begin{cases} Q_r = 0.5 \cdot rel_{LAI} \text{ if } \psi_x > -15,000 \\ \psi_x = -15,000 \text{ else} \end{cases}$,<br>$Q_r\vert _{tips} = 0$, no-flux at all soil faces | No reference solution, comparison amongst models | loam (Table 3), $k_x=0.0432$, $k_r=1.73 \times 10^{-4}$, RSA parameters from M1.1, $rel_{LAI}$ scales the potential transpiration |

<sup>1</sup> All paths are relative to the github repository https://github.com/RSA-benchmarks/collaborative-comparison.git. For other abbreviations and units see Table 1.

### How to participate

The participation includes three steps:

#### (1) Registration

Any interested researcher is welcome to contact the corresponding author of this paper, Andrea Schnepf, with the following information: Name, affiliation, name or reference to the participating simulator. Upon signing a letter of agreement confirming that results of other participants will not be published without consent, researchers will be accepted as participants and enabled to include their individual simulation results to the github repository of this benchmark initiative, https://github.com/RSA-benchmarks/collaborative-comparison.

#### (2) Simulation

Each participant implements all or a selected number of benchmark problems in their respective simulator and makes the results in the prescribed formats available to the github repository through pull requests. Requested formats include RSML (Lobet et al., 2015) for root architectures and VTK (Schroeder et al., 2006) for 3D and 1D simulation outputs. Python scripts to read and write RSML files will be provided on the github repository. Packages to read and write VTK files are for example available at https://pypi.org/project/vtk/.

#### (3) Analysis and publication

The analysis of results and computation of relevant metrics, such as root mean square error, coefficient of determination or Nash–Sutcliffe efficiency, will be done by the code implemented in the Jupyter Notebooks for each benchmark problem. The final goal is to jointly publish the results.

### 2.1 Benchmarks for individual modules

#### 2.1.1 Module M1: Root system architecture models

Root system architecture models (RSA models) are that module within a complex functional-structural plant model that simulates the structure, topology, and 3D placement of the roots. They simulate the growth of root systems as (upside down) tree-like structures based on rules regarding elongation, branching and death. Mostly, they are discrete models and represent the root system by a mathematical graph (i.e., nodes and edges/root segments). Each node or segment may be additionally associated with attributes such as radius, age or hydraulic properties.

The aim of this first benchmarking exercise is to determine if root architecture models currently available are able to reproduce realistic root architectures when being parameterised on the basis of a common experimental data set (Fig. 2a). The particular challenge to benchmark RSA models is to include the stochastic nature of these models. We propose to perform the benchmarking of those models in four steps: (1) Parameterising the root architecture models based on the provided experimental data, (2) Simulating a set of root systems for a dicotyledonous (*Lupinus albus*) and a monocotyledonous (*Zea mays*) plant species following two benchmark scenarios (M1.1 and M1.2), (3) Export and store the simulated root systems as Root System Markup Language (RSML) files (Lobet et al., 2015), and (4) Compare the simulation results using the data analysis pipelines available in the associated Jupyter Notebooks. The analysis pipelines are explained below and illustrated in Fig. 1. In particular, we include persistent homology as an approach that augments purely trait-based comparisons, i.e., two root systems with the same total root length could be very different based on the persistent homology approach.

**Figure 1:**
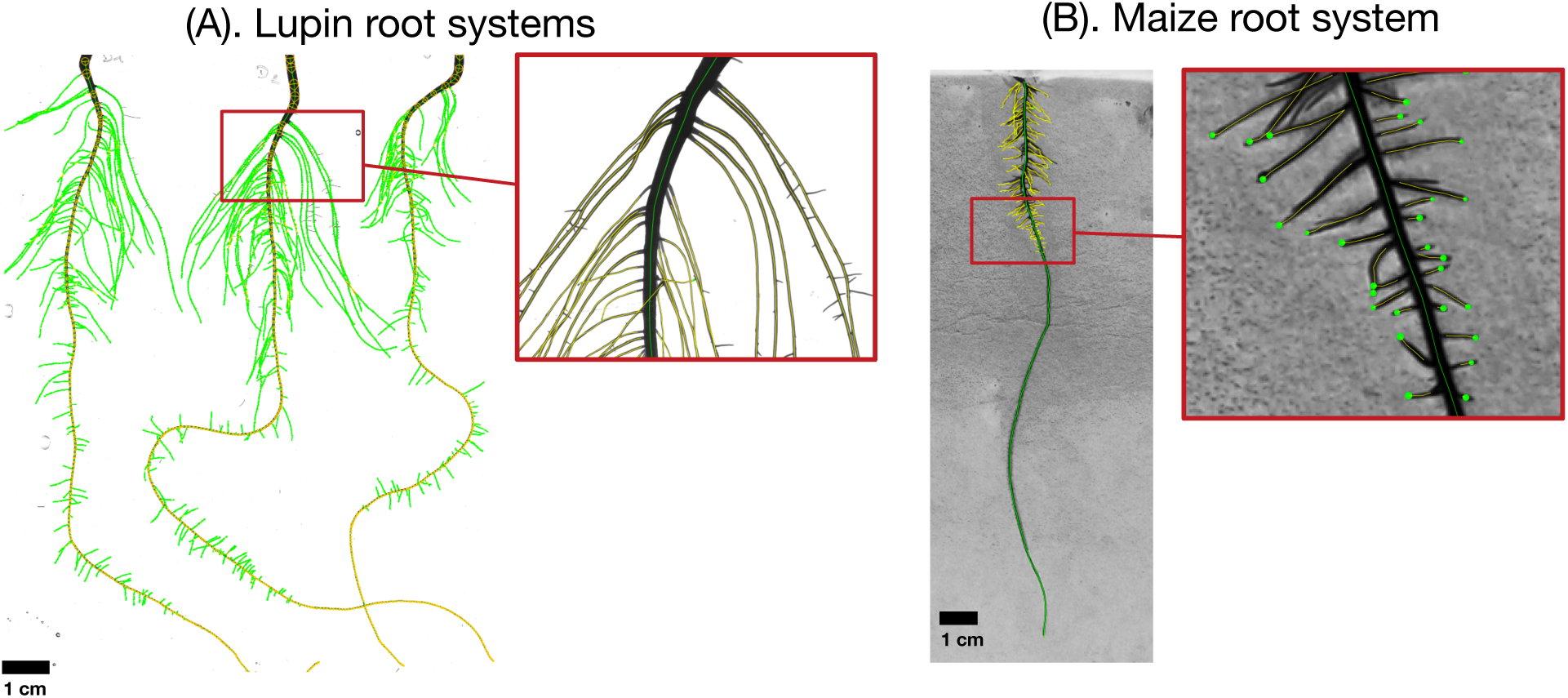
Example of root images used for the benchmarking dataset. Panel (A) shows an image of lupin root systems, 11 days old, growing in an aeroponic setup. Panel (B) shows an image of a maize root system growing on filter paper (5 days old). All images were analysed using the semi-automated root image analysis software SmartRoot (Lobet et al., 2011), colours distinguish different root orders. The RSML files containing the full information about the root systems are provided on the github repository in the folder “M1.1 RSA calibration\M1.1 Reference data”.

**Figure 2:**
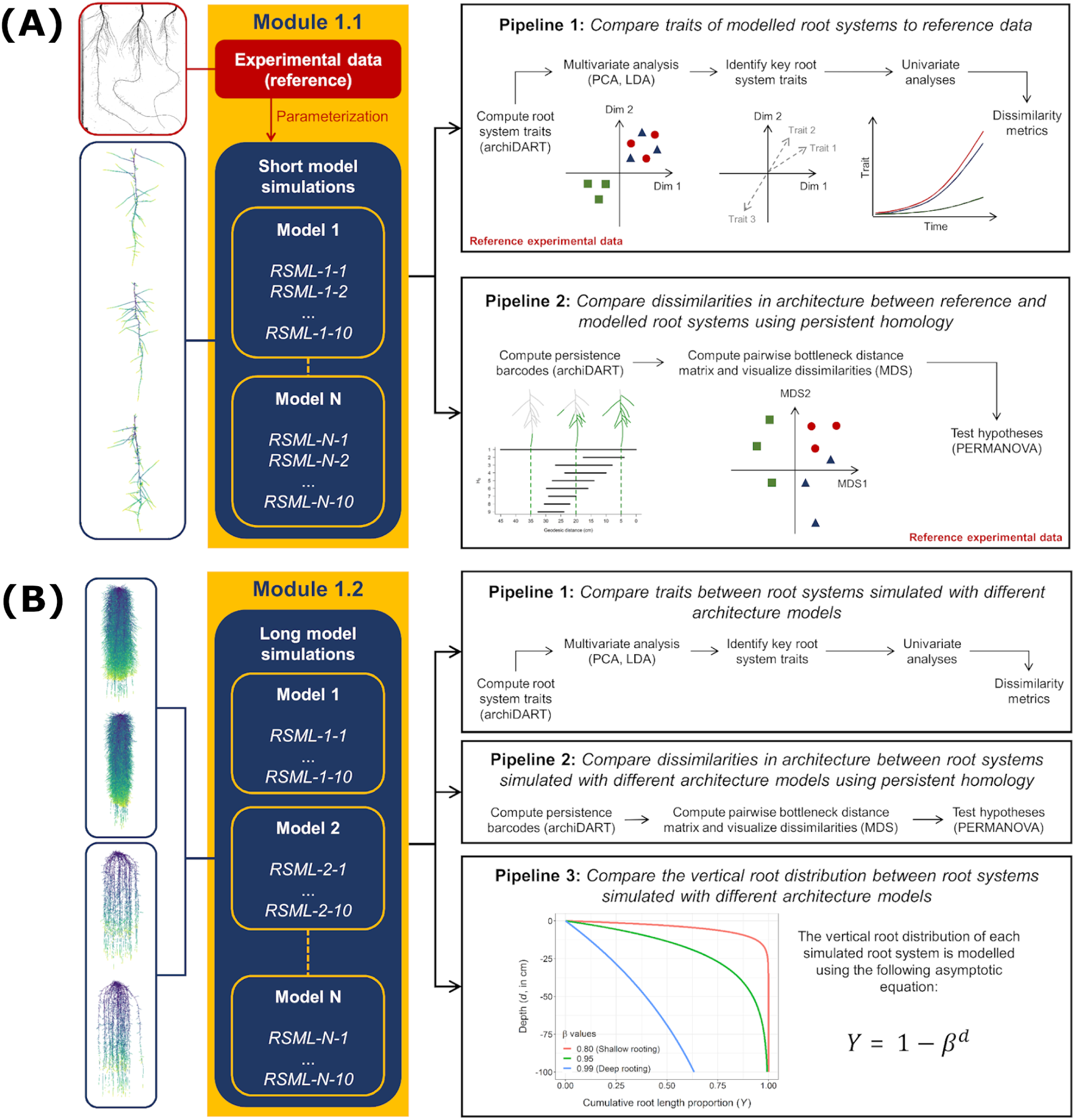
Presentation of the data analysis pipelines used for the benchmarking of root architecture models. Panels a and b show the first (M1.1) and second (M1.2) benchmark scenarios, respectively.

##### M1.1 Root system architecture model calibration

The different available root architecture models (see e.g. Dunbabin et al., 2013) are partly different in the way they represent the growth processes, i.e. we are looking at process-level differences between the different models. Thus, each participating RSA model will have a different set of parameters that drive root growth. This is the reason why, in this benchmark, we do not prescribe a parameter set as in e.g. M2 or M3, but we let each participating model derive its respective model parameters based on a reference dataset. In this first benchmark (M1.1), modellers simulate root systems for the same duration as the age of the root systems in the reference dataset.

###### Reference data set

Although the parameterisation of 3D models using a set of parameters derived from 2D images has some limitations, it has been shown to be a simple and efficient strategy allowing the simulation of realistic 3D root systems (Landl et al., 2018). Our reference dataset contains two distinct sets of images: (1) images of lupin roots grown for 11 days in an aeroponic setup (Lobet et al., 2011), and (2) images of maize roots grown for 8 days on filter papers (Hund et al., 2009). All images were analysed using the semi-automated root image analysis software SmartRoot (Lobet et al., 2011) and root tracings were saved as RSML files for further analysis (Fig. 1). These RSML files were then processed using functions of the R package archiDART developed to compute root system- and single root-level metrics (Delory et al., 2016, 2018). These metrics have been made open-access (https://github.com/RSA-benchmarks/collaborative-comparison/tree/master/root_architecture/data) and should help modellers to parameterise their respective RSA model.

###### Required output

The following results are to be uploaded via pull requests to this path on the github repository: M1 Root architecture development/M1.1 RSA calibration/M1.1 Numerical results.

1. A text file including the outcome of the calibration step, i.e., the set of model input parameters required for the specific simulator.
2. Simulation output from running the root architecture model using this parameter set in RSML format. Due to the stochastic nature of root architecture models, 100 realisations of each model setup are requested. The file format should be RSML and the file name should be of the form “modelname_replicate”, e.g. “CRootBox_1.rsml”.

###### Reference data analysis and automated model comparison

Statistical evaluation of a root architecture model has for example been done by Schnepf et al. (2018); Delory et al. (2018). This motivated the creation of two data analysis pipelines for the first benchmark (M1.1) that will be used to compare simulation outputs with reference experimental data (reference root systems) (Fig. 2a). These two data analysis pipelines are implemented in the Jupyter Notebook `RSA calibration.ipynb` that can be found on the github repository that contains code that will automatically include every model output in the analysis that is available in the prescribed folder. The analysis relies on the functions available in the **R** package archiDART (Delory et al., 2016, 2018). In the first pipeline, traits computed at the root system level (e.g., total root system length, number of roots per branching order) are compared between all simulated and reference root systems. This comparison takes place in three steps: (1) identifying the key morphological, architectural, and topological (Fitter indices, Fitter 1987; Fitter and Stickland 1991) traits explaining differences between simulated and reference root systems using multivariate data analysis techniques (e.g., discriminant analysis and principal component analysis), (2) looking at the point in time, beyond the time period for which there are measurements, when simulated and reference root systems start to diverge/converge with regard to the key root system traits identified in the previous step and how large these differences are, and (3) assess the degree of dissimilarity between simulated and reference root systems using dissimilarity metrics based on the raw data (Janssen and Heuberger, 1995).

In the second pipeline, dissimilarities in architecture between reference and simulated root systems are compared using persistent homology. Persistent homology is a topological framework that has proven to be a very powerful tool for capturing variations in plant morphology at different spatial scales (Li et al., 2017, 2018). The main output of a persistent homology analysis is a persistence barcode recording the appearance and disappearance of each root branch when a distance function traverses the branching structure (see Fig. 1 in Delory et al., 2018). The degree of similarity between different root system topologies can be assessed by computing a pair-wise distance matrix to compare persistence barcodes. In addition, Delory et al. (2018) showed that both trait-based and persistent homology approaches nicely complement each other and allow root researchers to more accurately describe differences in root system architecture (Delory et al., 2018). In our data analysis pipeline, a persistent homology analysis comprises the following steps: (1) computing a persistence barcode for each simulated and reference root system using a geodesic distance function, (2) computing dissimilarities between persistence barcodes using a bottleneck distance, (3) visualize dissimilarities between root systems using multidimensional scaling, and (4) test specific hypotheses using permutational multivariate analysis of variance (PERMANOVA) (Anderson, 2001).

##### M1.2 Long model simulations

In this benchmark, modellers use the same input parameter set as in M1.1, but simulate root system growth and development for a longer time period (60 days). The aim of this second benchmarking exercise is to assess if the different models diverge (or converge) if simulations are run for a longer time period and extrapolate beyond the provided data set (Fig. 2b). This is of great importance, as parameterisation of RSA models is often based on relatively young plants, whereas knowledge of RSA of older root systems is scarce. Therefore, for this M1.2 scenario, experimental data are not used as the basis of comparison anymore. It has to be noted that these two benchmark problems focus on root architecture dynamics modelling only, thus effect of soil properties on root growth is not explicitly modelled.

###### Required output

The following results are to be uploaded via pull requests to this path on the github repository: `M1 Root architecture development/M1.2 RSA simulation/M1.2 Numerical results.`

1. A text file including the model input parameters used for the specific simulator.
2. Simulation output from running the root architecture model using this paramter set in RSML format. Due to the stochastic nature of root architecture models, 100 realisations of each model setup are requested. The file file format should be RSML and the file name should be of the form “modelname_replicate”, e.g. “CRootBox_1.rsml”.

###### Analysis pipeline for M1.2

For the second benchmark (M1.2), three data analysis pipelines are used to compare simulation outputs given by different root architecture models. For this benchmark, the reference experimental data cannot be used as a reference as data of 60 day old plants is not available. The first two data analysis pipelines for M1.2 are very similar to the ones described earlier for the M1.1 benchmark. First, model outputs are compared using morphological, architectural, and topological traits computed at the root system level. Second, differences in root system morphology are analysed using persistent homology. In addition to these two analysis pipelines, we included a third one to analyse differences in vertical root distribution between root systems simulated with different root architecture models. To do so, we use the modelling approach described in Oram et al. (2018). Briefly, relative cumulative root length density (Y(d)) is computed using Eq. (1)

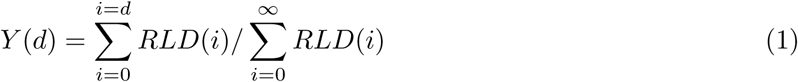

Eq. (2) is fitted to the computed Y(d) using a non-linear least square means fitting procedure. The fitting constant *β* is used to compare modeled rooting depth, with high *β* corresponding to deep rooting.

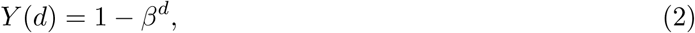

#### 2.1.2 Module 2: Water flow in soil only

In this module, we describe benchmark problems that only relate to water flow in soil. Water flow in soil is most commonly described by the Richards equation in three dimensions:

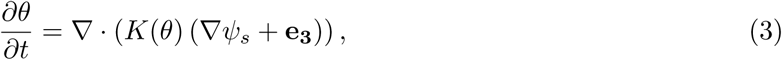

where *θ* is the volumetric soil water content (cm^3^cm^−3^), *K* is the hydraulic conductivity (cm day^−1^), *ψ*_*s*_ is the soil water pressure head (cm), and **e**_**3**_ = (0, 0, 1) is the downward unit vector.

The relationship between soil water pressure head and water content is generally described by the water retention curve. In the following we will use the van Genuchten equation (Van Genuchten, 1980) to describe this curve specifying the soil moisture characteristic of specific soils.

We expect differences between the outputs of different simulators to be mainly numerical solution-level differences, i.e., due to numerical scheme and implementation. Different numerical solutions of the Richards equation have been analysed before, and for some settings analytic solutions exist. We will use the benchmarks presented by Vanderborght et al. (2005) to benchmark the part of the participating functional structural root architecture models where water movement in soil is described. The analytical solutions provided in that paper are related to vertical changes in the soil profile only. As most functional-structural root architecture models have a 3D soil module, they will prescribe no-flux boundary conditions at the sides of a domain with 25 cm length and width for the numerical implementation of those problems.

In the following we will describe the benchmarks for water movement in soil. Table 3 gives an overview of the soil hydraulic properties that will be used throughout all the benchmarks involving water flow in soil.

**Table 3:** Soil hydraulic taken from Vanderborght et al. (2005). *θ*_*res*_ is the residual water content, *θ*_*sat*_ is the saturated water content, *α* and *n* are the van Genuchten parameters, *K*_*sat*_ is the saturated soil hydraulic conductivity and *λ* is the van Genuchten-Mualem parameter

| Soil type | $\theta_{res}$<br>(-) | $\theta_{sat}$<br>(-) | $\alpha$<br>(cm<sup>-1</sup>) | $n$<br>(-) | $K_s$<br>(cm d<sup>-1</sup>) | $\lambda$<br>(-) |
| --- | --- | --- | --- | --- | --- | --- |
| sand | 0.045 | 0.43 | 0.15 | 3.0 | 1000 | 0.5 |
| loam | 0.08 | 0.43 | 0.04 | 1.6 | 50 | 0.5 |
| clay | 0.1 | 0.40 | 0.01 | 1.1 | 10 | 0.5 |

##### M2.1: Infiltration

This benchmark scenario is taken from Vanderborght et al. (2005). All parameters, initial and boundary conditions are given in Table 2 and are described below. For each of the soil types, sand, loam and clay, we consider the rate of infiltration into a soil with an initial homogeneous soil water pressure head of *ψ*_*s*_ =-400 cm. All profiles are 200 cm deep, at the top boundary we prescribe a constant influx of 100 cm d^−1^ as long as the soil is still unsaturated, and a Dirichlet boundary condition of *ψ*_*s*_=0 cm as soon as the soil is fully saturated. At the bottom boundary, we prescribe free drainage. Since this problem only produces gradients in the vertical direction, we compare numerical model results with the 1D analytical solution described in Vanderborght et al. (2005).

###### Reference solution

The analytical solution is given by the travelling wave equation

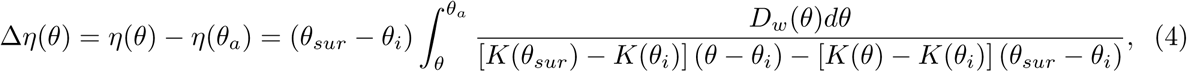

where *D*_*w*_ is the water diffusivity (defined as 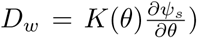, *θ*_*sur*_ is the water content at the soil surface, *θ*_*i*_ is the initial water content, *θ*_*a*_ is a reference water content (taken to be *θ*_*a*_ = (*θ*_*sur*_ + *θ*_*i*_)/2), 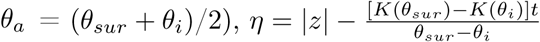 and Δ*η*(*θ*) is the distance of the front to the the position of the reference water content. The implementation of this analytical solution, implemented in the Jupyter Notebook `M2.1 Benchmark problem.ipynb`, reproduces Figure 4abc from Vanderborght et al. (2005), where the water content is plotted after 0.1, 0.2, and 0.3 days for the sand scenario; 0.2, 0.5, and 1 days for the loam scenario; and 0.1, 0.2, and 0.5 days for the clay scenario (see Fig. (3)).

**Figure 3:**
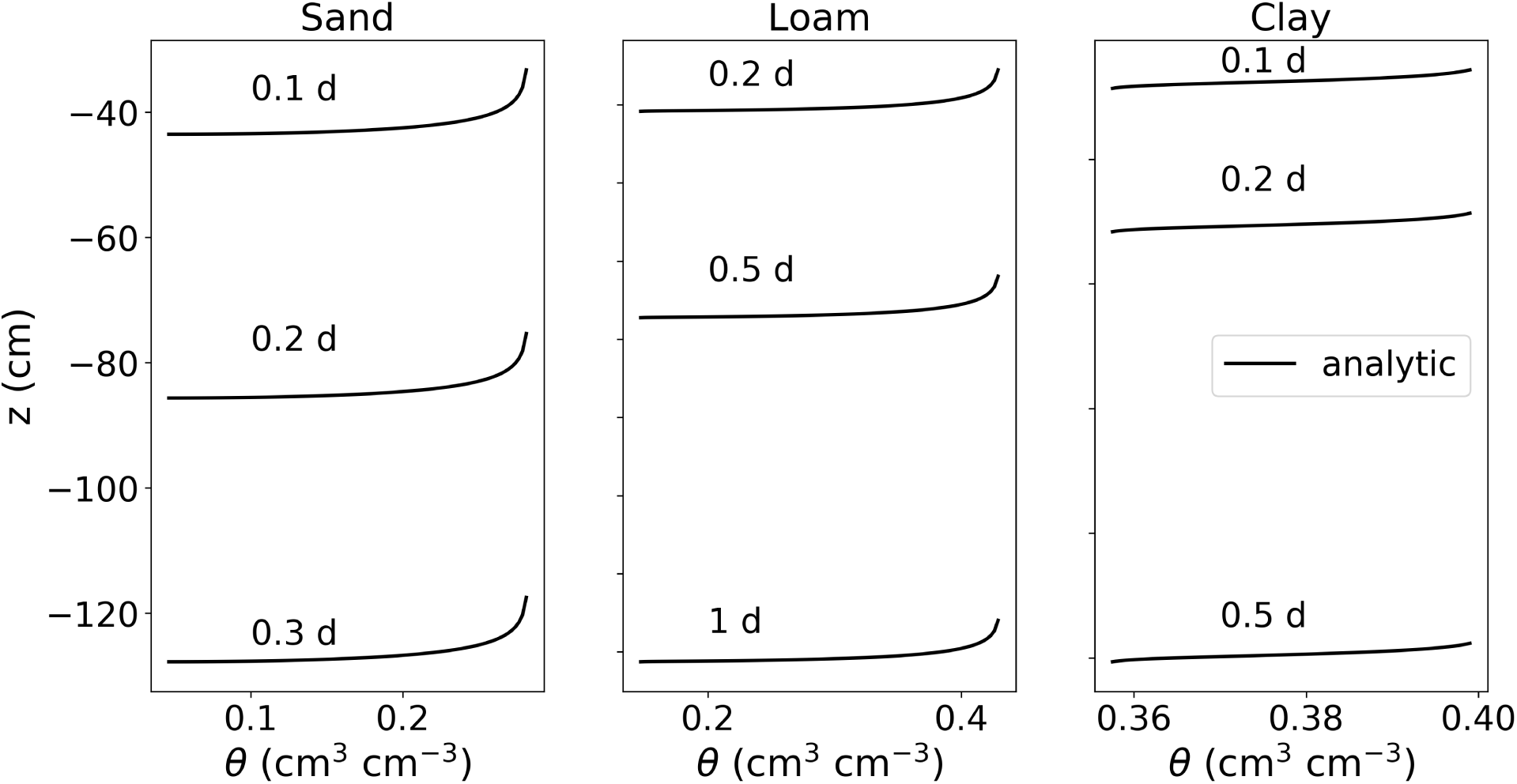
Results of M2.1: Infiltration into three initially dry soils: sand, loam and clay.

**Figure 4:**
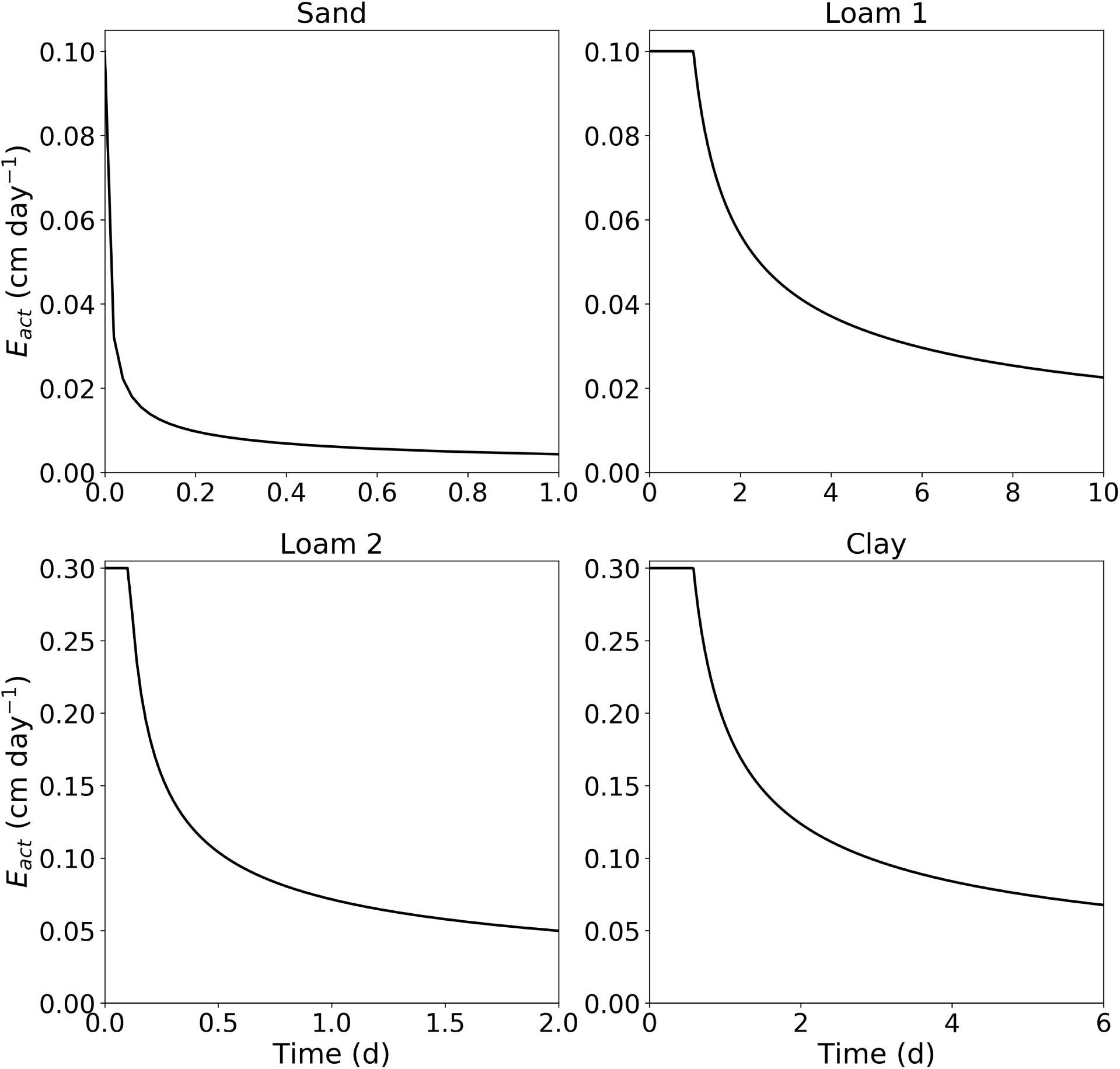
Results of M2.2: Rate of evaporation with respect to time from sand with *J*_*s,pot*_=0 1cm/d, loam with *J*_*s,pot*_=0 1cm/d, loam with *J*_*s,pot*_=0 3cm/d and clay with *J*_*s,pot*_=0 3cm/d

###### Required output

The following simulation results of participating models are to be uploaded via pull requests to this path on the github repository: `M2 Water flow in soil/M2.1 Infiltration/M2.1 Numerical results.`

1. A text file consisting of two rows containing comma separated depth values (cm) in the first, and water content (cm^3^cm^−3^) in the second for each time point and infiltration scenario (i.e. 3 (time points) × 3 (scenarios) results = 18 rows). The file name should be of the form “simulatorname.txt”, e.g. “DuMux.txt”.

Note that we do not prescribe spatial or temporal resolution of the outputs, as that may depend on the individual numerical schemes.

##### M2.2: Evaporation

This benchmark reproduces Fig. (5) of Vanderborght et al. (2005). We consider four scenarios (sand, loam 1, loam 2, clay) in which we are interested in the actual evaporation over time from an initially moist soil (*ψ*_*i*_ = − 40cm for the sand scenario and *ψ*_*I*_ = − 200cm for all other scenarios). The domain is 100 cm deep with a width and length of 10 cm. At the top boundary, we prescribe a constant efflux of *J*_*s,pot*_=0.1 cm d^−1^ for the sand and loam 1 scenario, and 0.3 cm/day for the loam 2 and clay scenarios, at the bottom we prescribe zero-flux. When the soil reaches a critical soil water pressure head of −10.000 cm at the surface, we switch to a Dirichlet boundary condition with *ψ*_*s*_= −10.000 cm.

**Figure 5:**
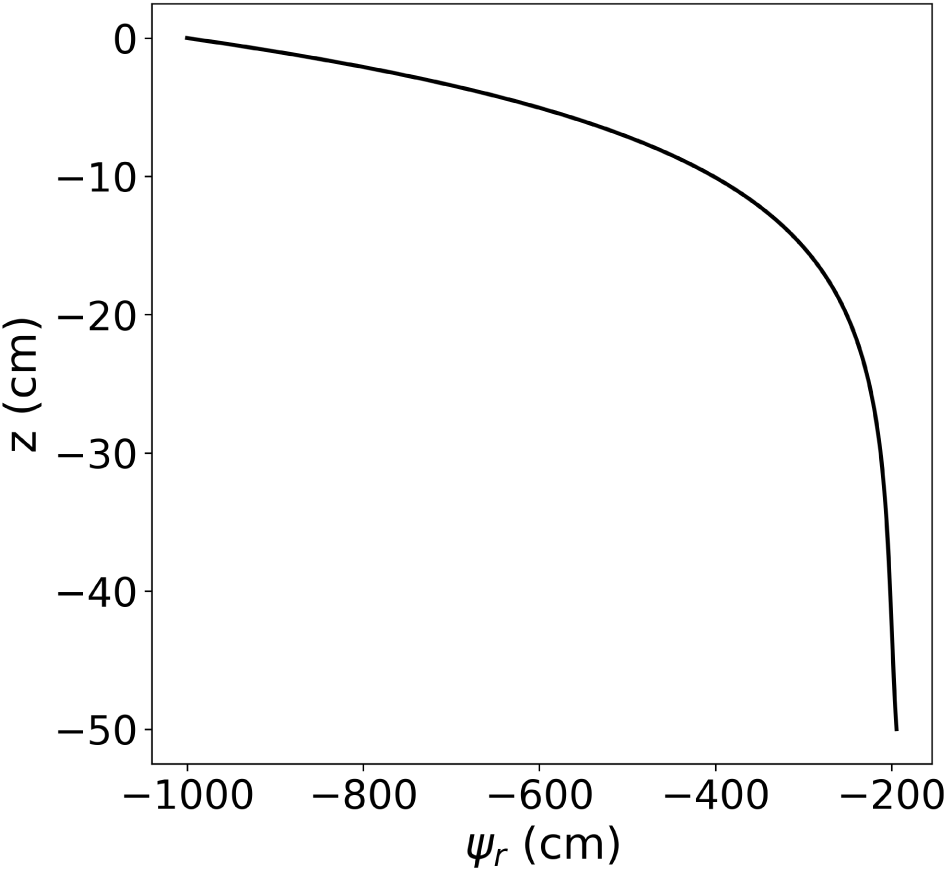
Results of M3.1: Root water pressure head distribution within a single vertical root.

###### Reference solution

The analytical solution to this problem is given by

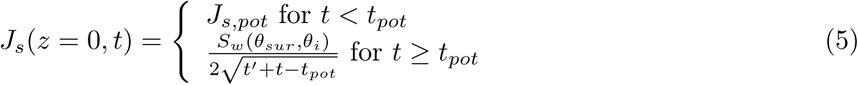

where 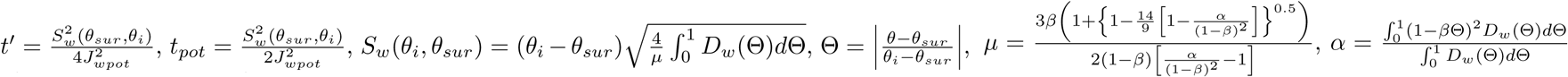, and 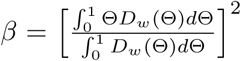. Fig. (4) shows the rate of evaporation over time for the four scenarios soil, loam 1, loam 2, clay.

###### Required output

The following simulation results of participating models are to be uploaded via pull requests to this path on the github repository: `M2 Water flow in soil/M2.2 Evaporation/M2.2 Numerical results.`

1. A text file consisting of two rows containing comma separated depth values (cm) in the first, and root pressure head (cm) in the second for each scenario (i.e. 4 (scenarios) × 2 (rows) = 8 rows). The file name should be of the form “simulatorname.txt”, e.g. “DuMux.txt”.

Note that we do not prescribe spatial or temporal resolution of the outputs, as that may depend on the individual numerical schemes. It is the responsibility of each participant, to upload the best possible solution.

#### 2.1.3 Module 3: Water flow in roots

In this benchmark, we consider water flow in xylem with constant and homogeneous soil water pressure head. This problem is well described, e.g., in Roose and Fowler (2004) and Doussan et al. (1998). Its analytical solution for a single root was already derived by Landsberg and Fowkes (1978). In Appendix A, we present a derivation that is equivalent to the solution of Landsberg and Fowkes (1978) but uses exponential instead of hyperbolic functions. Briefly, conservation of mass in a branched root network with both axial and radial water flow, neglecting plant water storage and osmotic potential, yields Eq. (6),

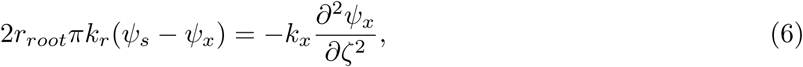

where *r*_*root*_ is the root radius (cm), *k*_*r*_ is the radial conductivity (d^−1^), *ψ*_*s*_ is the soil water pressure head of the surrounding soil (cm), *ψ*_*x*_ is the root water pressure head inside the xylem (cm), *k*_*x*_ is the axial conductance (cm^3^ d^−1^), and *ζ* is the axial coordinate (cm).

##### M3.1: A single root in static soil with constant root hydraulic properties

In this benchmark problem, we assume a vertical single straight root segment surrounded by a soil with a constant and uniform soil water pressure head (i.e. the soil is not in hydrostatic equilibrium). We prescribe the root water pressure head at the root collar as *ψ*_*x*_ |_collar_ = *ψ*_0_, and no axial flow at the root tips.

###### Reference solution

For constant *k*_*r*_ and *k*_*x*_ we can solve Eq. (6) yielding

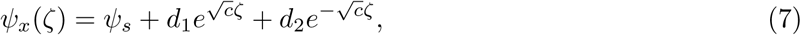

with *c* = 2*r*_*root*_*πk*_*r*_*/k*_*x*_. The integration constants *d*_1_ and *d*_2_ for above boundary conditions are given by

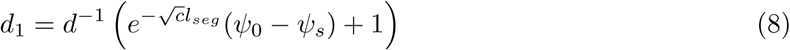

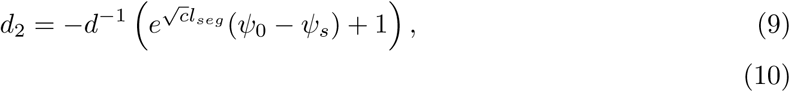

where *l*_*seg*_ is the segment length, and *d* is the determinant of above matrix

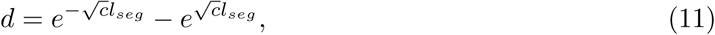

see Appendix A. Fig. 5 shows the analytical solution to this benchmark using the parameters given in Table 4.

**Table 4:** Parameters of scenario M3.1.

|  |  |  |
| --- | --- | --- |
| l | 50 | length of a single straight root (cm) |
| $r_{\text{root}}$ | 0.02 | radius (cm) |
| $k_z$ | $4.32 \times 10^{-2}$ | axial conductivity ($\text{cm}^3 \text{d}^{-1}$) |
| $k_r$ | $1.73 \times 10^{-4}$ | radial conductivity ($\text{d}^{-1}$) |
| $\psi_s$ | -200 | static soil water pressure head (cm) |
| $\psi_0$ | -1,000 | Dirichlet boundary conditions at the root collar (cm) |

###### Required output

The following simulation results of participating models are to be uploaded via pull requests to this path on the github repository: `M3 Water flow in roots/M3.1 Single root/M31 Numerical results/.`

1. A text file consisting of two rows containing comma separated depth values (cm) in the first, and root pressure head (cm) in the second. The file name should be of the form “simulatorname.txt”, e.g. “DuMux.txt”.

Note that we do not prescribe spatial resolution of the outputs, as that may depend on the individual numerical schemes.

##### Benchmark M3.2: A small root system in a static soil

In the following benchmark, we extend benchmark M3.1 from a single root to a root system. We consider water flow inside a small static root system of a lupine plant which was grown for 14 days in a soil-filled column of 20 cm depth and 7 cm diameter. The root system was imaged by MRI at Forschungszentrum Jülich; the segmented root structure is provided in RSML, DGF (Dune grid format) (Bastian et al., 2008) and RSWMS (Javaux et al., 2008) formats in the folder `M3 Water flow in roots/M3.2 Root system/root_grid` on the github repository. It is visualised in Fig. 6(a,b) with colours denoting root order and root segment age.

**Figure 6:**
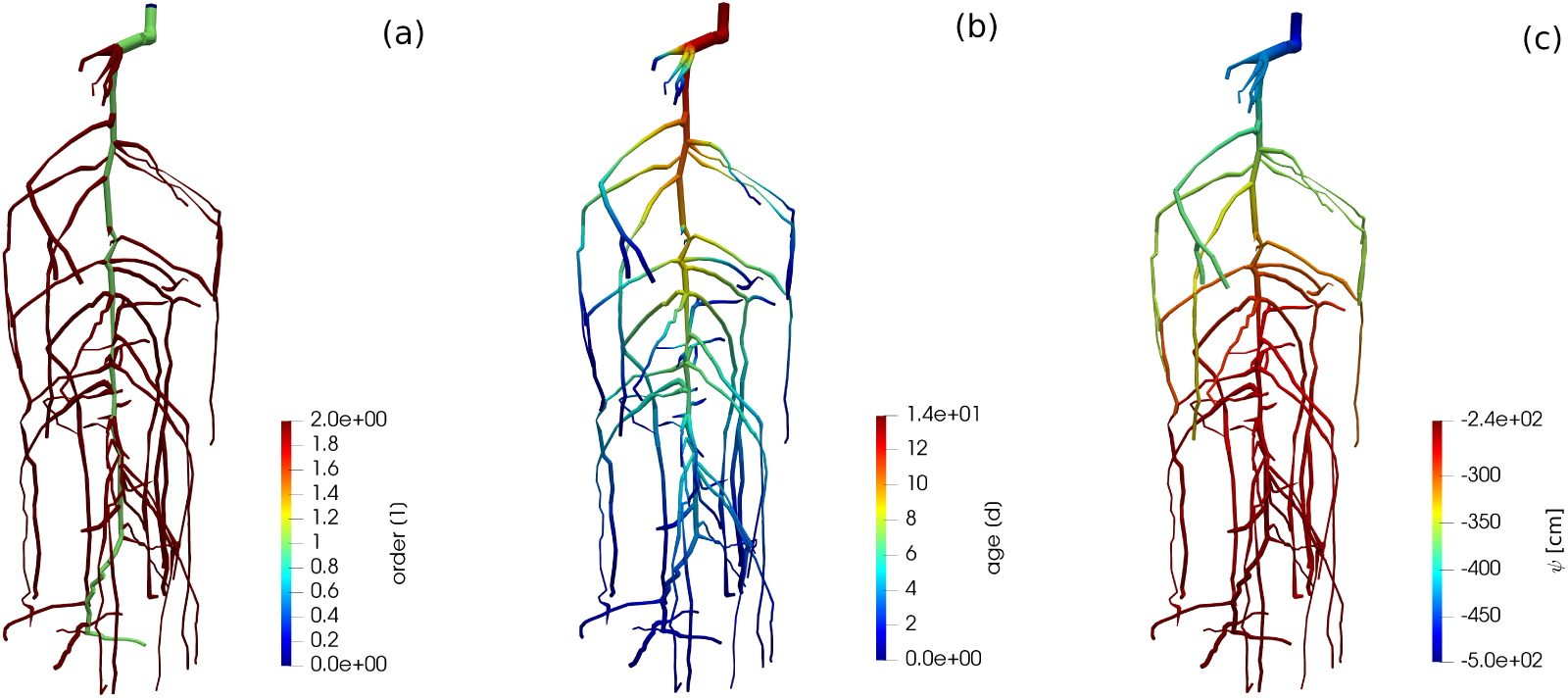
Visualisation of the root system of M3.2 with colours denoting (a) root order, (b) root segment age, (c) root water pressure head.

###### Reference solution

The reference solution for this problem is given by the hybrid analytical-numerical solution of water flow in the root hydraulic architecture proposed by Meunier et al. (2017). The advantage of this solution is that it is independent of the spatial resolution of the root system (i.e. root segment length).

We consider two scenarios. The first one uses the same constant root hydraulic properties as given in Table 4, i.e. considering the same root hydraulic properties for each root segment. In the second scenario, we consider age-dependent root hydraulic properties for tap root and laterals of lupine as obtained by Zarebanadkouki et al. (2016, exponential function scenario) and converting distance from root tip to root age by assuming a root growth rate of 1 cm d^−1^. This parameterisation takes into account that roots get a higher axial conductivity and lower radial conductivity as they are becoming older (see Fig. 7, a table with the actual values is provided on the github repository, in: `M3 Water flow in roots/M3.2 Root system/M3.2 Benchmark problem. ipynb.`

**Figure 7:**
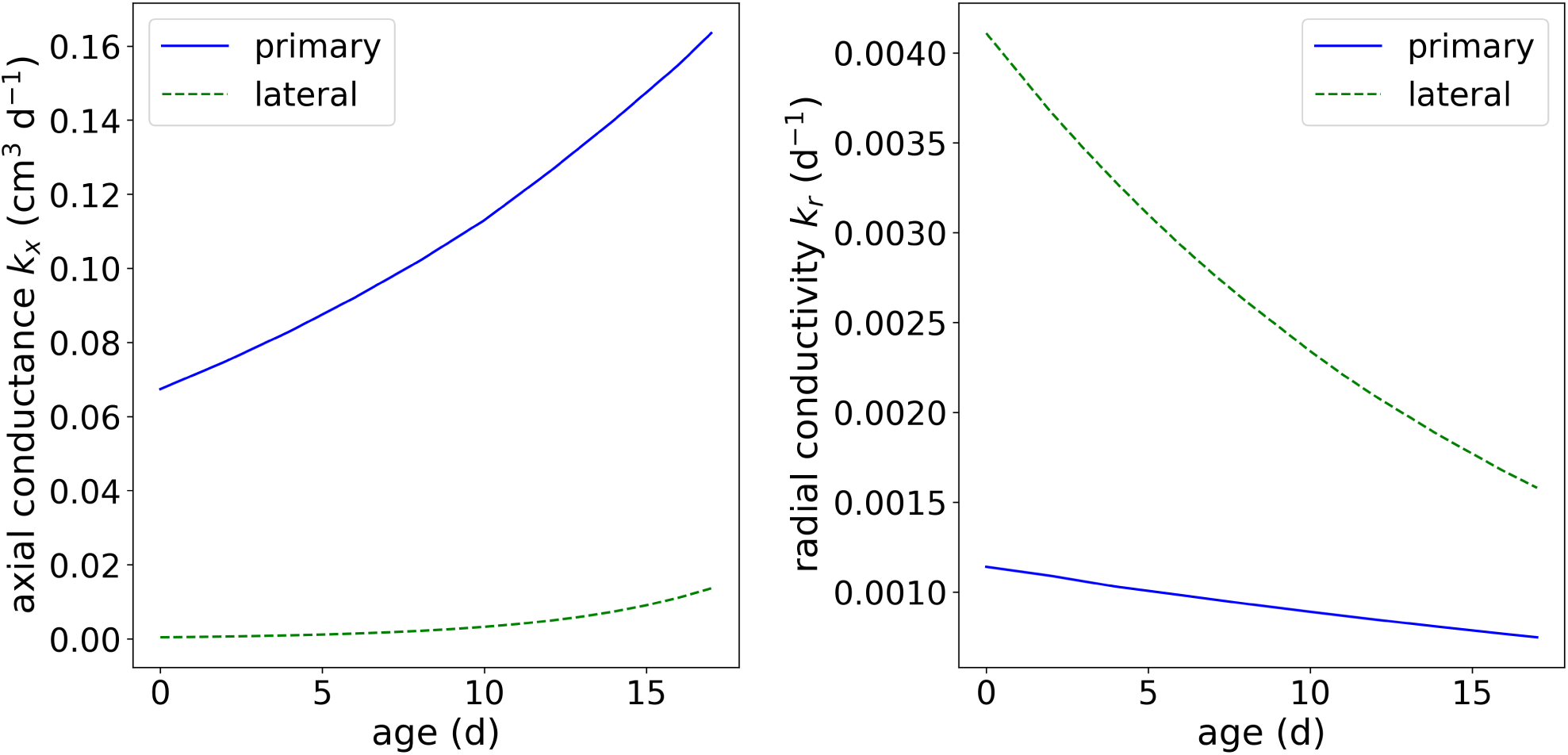
Root hydraulic properties dependency on root type and root segment age.

A sample 3-D visualisation of the model output is shown in Fig. 6(c) for the constant root hydraulic properties scenario. Fig. 8 shows the effect of constant and age-dependent root hydraulic properties under otherwise same (soil and boundary) conditions.

**Figure 8:**
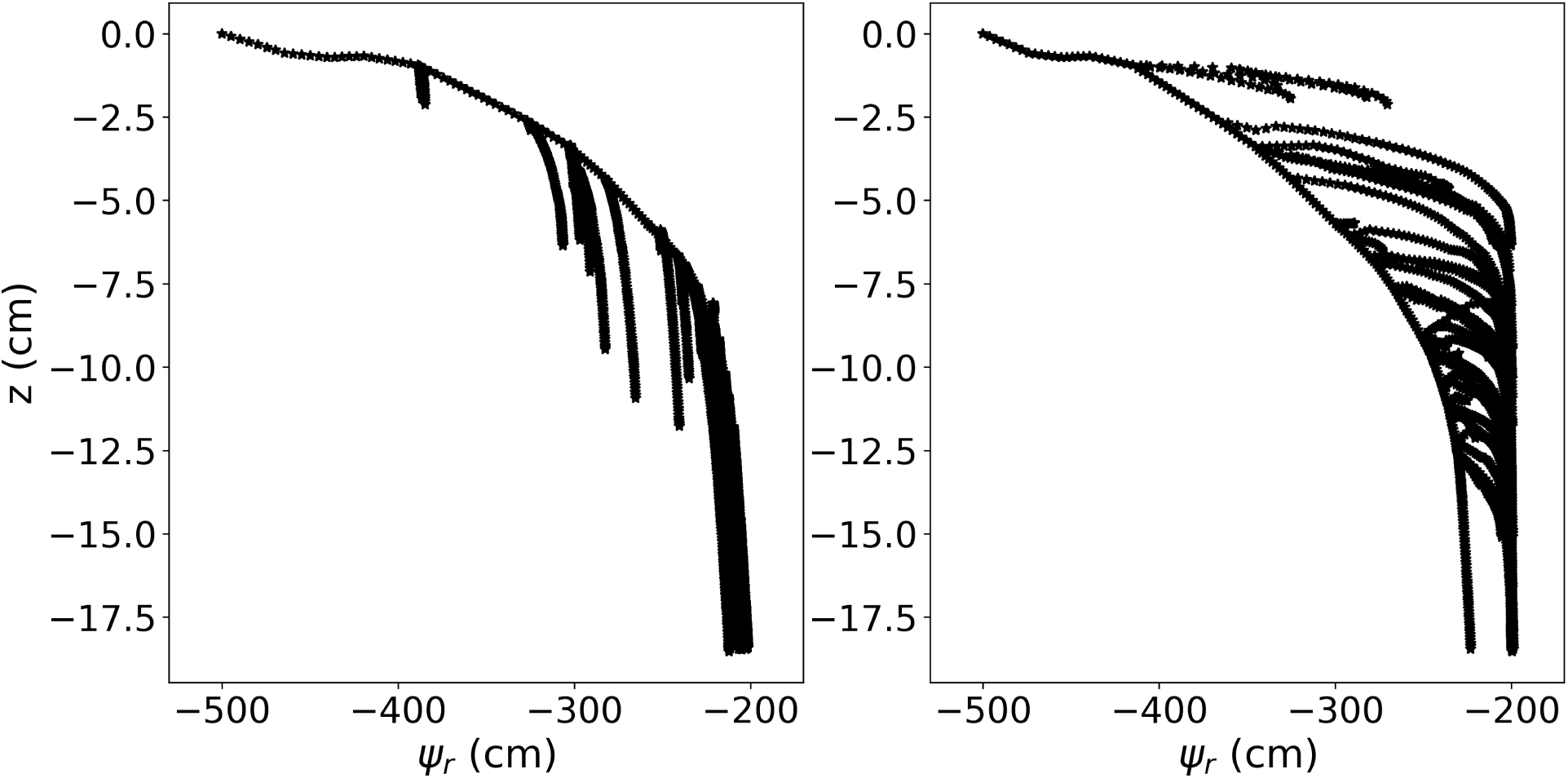
Results of M3.2. Left panel: Xylem pressure in each root segment of a root system with constant hydraulic properties. Right panel: Xylem pressure in each root segment of a root system with age-dependent hydraulic properties.

###### Required output

The following simulation results of participating models are to be uploaded via pull requests to this path on the github repository: `M3 Water flow in roots/M3.2 Root system/M32a Numerical results and M3 Water flow in roots/M3.2 Root system/M32b Numerical results` for the constant and age-dependent root hydraulic properties cases.

1. A text file consisting of two rows containing comma separated depth values (cm) in the first, and root pressure head (cm) in the second. The file name should be of the form “simulatorname.txt”, e.g. “DuMux.txt”.

Note that we do not prescribe spatial resolution of the outputs, as that may depend on the individual numerical schemes.

#### 2.1.4 Coupled benchmark scenarios C1: Root water uptake by a static root system

The way of coupling can easily introduce differences in simulated results because of numerical errors (especially when there is two way coupling) or because different assumption are made when implementing the coupling. No analytical solutions exists for the coupled problems presented here, but the coupling (C) benchmarks are intended to quantify differences between model outputs of coupled models. We may see differences observed in the non-coupled benchmarks to be amplified, or to be irrelevant for the coupled problem.

##### C1.1: Water uptake by a single root

This benchmark follows the paper of Schröder et al. (2008). Here we aim to see to what extent the different participating models can reproduce the hydraulic conductivity drop near the root surface under different soil conditions and transpiration demands. Thus, it requires the participating line-source based models to strongly increase the spatial resolution of the 3D soil domain. From this benchmark, we will learn, whether the spatial resolution required to reproduce radial soil water pressure head gradients would be in a feasible order of magnitude for larger soil-root systems or not. If not, there are approaches to estimate soil water pressure head drop at the root-soil interface from bulk soil values as e.g. in Mai et al. (2019); Beudez et al. (2013), see also benchmark C1.2.

###### Reference solution

The analytical solution is based on the analytical solutions of the 1D radially symmetric problem of water uptake by a single root, in which root water uptake is described as a boundary condition at the root-soil interface. We consider here two water uptake regimes, a non-stressed condition with maximum root uptake (*q*_*root*_), and a stressed condition with a limiting plant root water potential constraining uptake. Based on the steady-rate assumption and using the matric flux potential 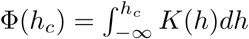 that linearises the Richards equation, the radial soil water pressure head profiles for non-stressed and stressed conditions (stress conditions are given when the soil water pressure head at the root surface reaches −15 000cm) are given by

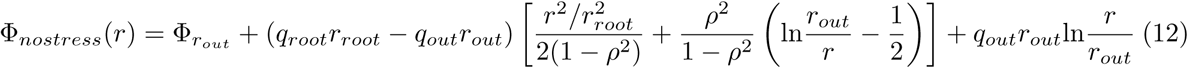

and

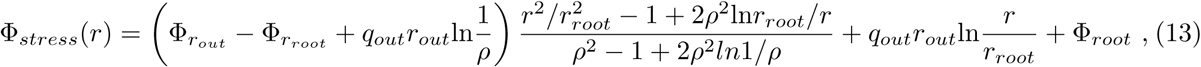

where 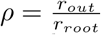.

Given the soil water pressure head at the outer boundary, the solution computes the soil water pressure head profile towards the root. Due to the steady-rate assumption, the problem has become a stationary boundary value problem. However, under non-stressed conditions, we can calculate the time that corresponds to a given radial soil water pressure head profile by dividing the volume of water removed from the soil domain by the known water flow rate. The water remaining in a 1 cm long hollow cylinder around the root is given by

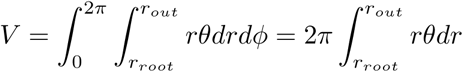

*θ* being the water content. The initially available water volume in the soil domain is given by

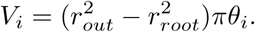

Thus, until the onset of stress, the corresponding time at which a given radial profile is reached is given by

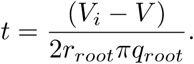

For the three soils sand, loam, and clay (Table 3), we compute the analytical solution with the following parameters: *r*_*root*_= 0 02cm, *r*_*out*_ = 1cm, *q*_*root*_ = 0 5cm/d, *ψ*_*s,lim*_ = −15000cm, *q*_*out*_ = 0cm/d and for different soil water pressure heads at the outer end of the cylinder. Fig. 9 shows the soil water pressure head gradients at the onset of stress (i.e., when the soil water pressure head at the root surface reached −15 000cm) and the time of its occurrence. The value of the initial water content is taken to be *θ*_*i*_ = −100cm. This analytical solution is for radial water flow in soil towards the root only, i.e., not considering gravity or water flow inside the roots. Ideally, in their numerical implementation of this benchmark, the different participating models will turn off gravity effects. The soil domain for this numerical implementation has a size of *l* × *w* × *d* = 1× 1 × 1 cm. The horizontal spatial resolution is high enough such that hydraulic conductivity drop near root surface can be resolved. The axial and radial conductances are high, such that the pressure inside the root is everywhere the same and the uptake flux is uniform.

**Figure 9:**
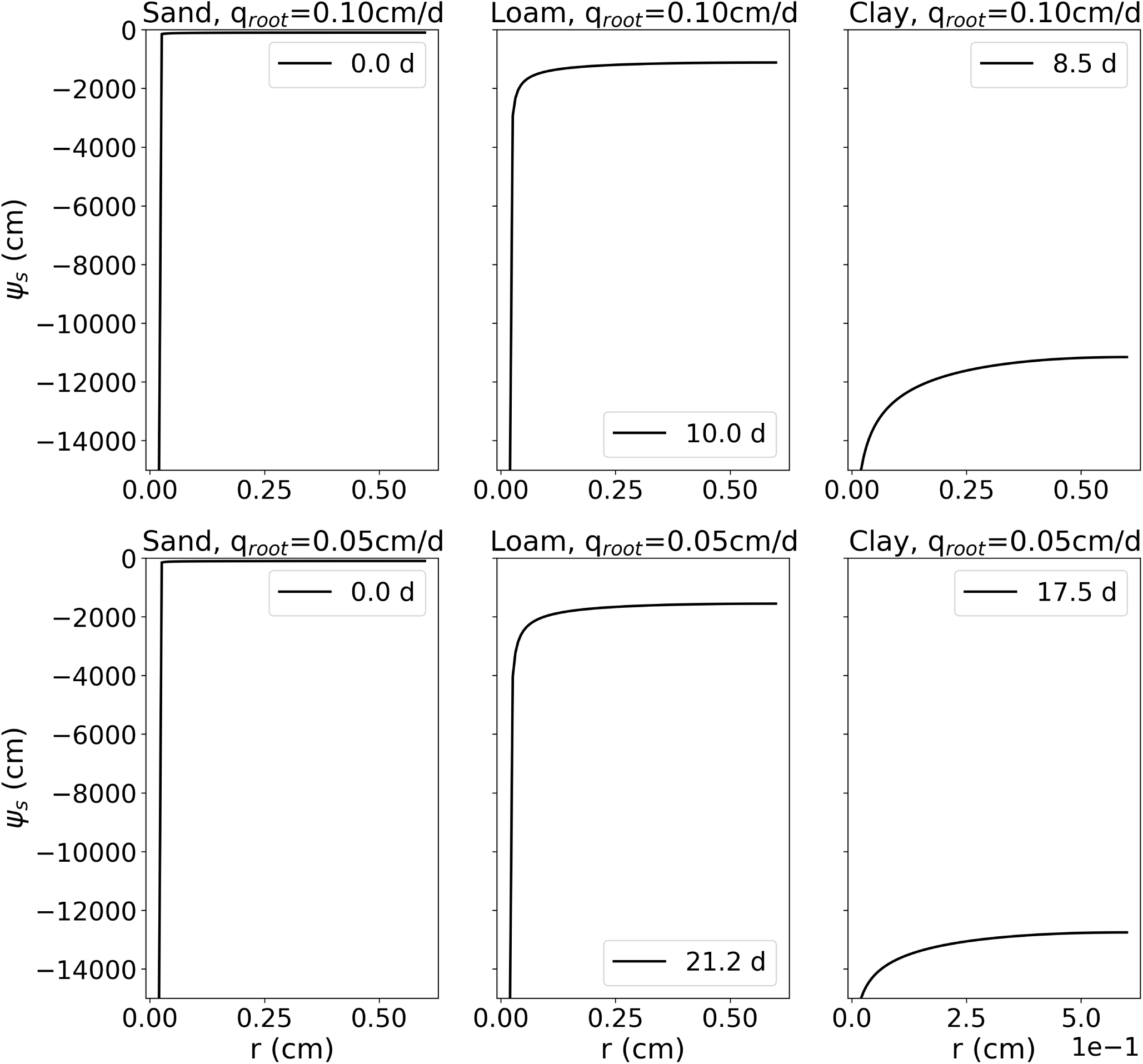
Results of C1.1: Soil water pressure head gradients around a single, transpiring, root at the onset of stress and the time of its occurrence

###### Required output

The following simulation results of participating models are to be uploaded via pull requests to this path on the github repository: `M3 Water flow in roots/M3.2 Root system/M32a Numerical results and M3 Water flow in roots/M3.2 Root system/M32b Numerical results` for the constant and age-dependent root hydraulic properties cases.

1. A text file consisting of two rows containing comma separated radial distances from the root surface (cm) in the first, and soil pressure head (cm) in the second for each soil and transpiration rate scenario (i.e., 3 (soils) × 2 (transpiration rates) × 2 = 12 rows. The file name should be of the form “simulatorname.txt”, e.g. “DuMux.txt”.

Note that we do not prescribe spatial or temporal resolution of the outputs, as that may depend on the individual numerical schemes.

#### 2.1.5 C1.2: Water uptake by a root system from drying soil

This benchmark scenario considers water uptake by a static 8-day-old lupine root system given in the public data set (Koch, 2019) as RSML or DGF. The root is the same as the one in benchmark M3.2, only younger, in order to reduce the computational cost for the reference scenario. The root system has been segmented from MRI measurements. The lupine is embedded in a soil box of *l* × *w* × *d* = 8 × 8 × 15 cm filled with loam (soil hydraulic properties given in Table 3). The benchmark is to evaluate the accuracy of root water uptake models under conditions of drying soil. To this end, the soil has an initial water content of *θ*_top_ = 0.129, corresponding to a pressure head *ψ*_*s*,top_ = −659.8 cm at the soil surface (*z* = 0). The pressure head in the rest of the domain initially follows a hydrostatic distribution

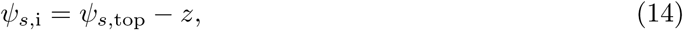

where *z* (in cm) denotes the vertical position (upward-pointing axis, zero at soil surface). At all soil boundaries, as well as at the root tips, no-flux boundaries are prescribed. A potential transpiration rate is given as the sinusoidal diurnal function

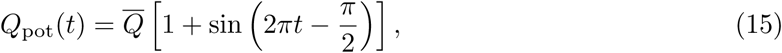

where the mean transpiration rate is 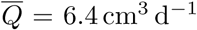, the time *t* is given in days, and *Q*_pot_(*t* = 0) = 0, that is, the simulation starts at night. The potential transpiration rate *Q*_pot_, Eq. (15), is enforced at the root collar (Neumann boundary condition) as long as the root water pressure head at the root collar is above *ψ*_*x*,crit_ = − 15 290 cm (corresponding to − 1.5 MPa). If this critical root water pressure head at the root collar is reached, the boundary condition is switched to a Dirichlet type boundary condition, enforcing a constant pressure head *ψ*_*x*,crit_ = − 15.290 cm at the root collar. This informal description is intentional, as the actual implementation of such a boundary condition may vary from simulator to simulator. We consider two scenarios. In scenario C1.2a the root hydraulic properties are constant. The tap root and lateral root conductivities are *k*_*x*_ = 4.32 × 10^−2^ cm^3^ d^−1^ and *k*_*r*_ = 1.73 × 10^−4^ d^−1^ (Table 4). For scenario C1.2b the root hydraulic properties depend on the root type and root age and are shown in Fig. 7.

Given the soil domain Ω and the network of root center-lines Λ, we solve the following coupled system of equations

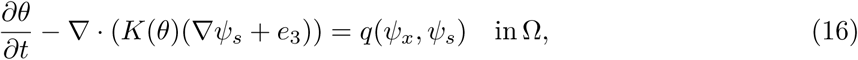

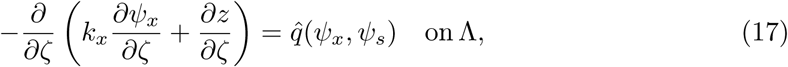

subject to the boundary conditions specified above, where *ζ* is a scalar parametrisation (local axial coordinate) of the root segments. The specific radial flux 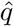 in units (cm^2^ d^−1^) is given by the average soil water pressure head on the root surface. The formulation of *q* in Eq. (16) may be different between different participating models. A discussion on singularity issues when evaluating the soil water content at the root center line can be found in Koch et al. (2018b). In many cases, the soil discretisation is much larger than the root diameter, and thus the drop in hydraulic conductivity near the root surface in dry soils may not be sufficiently resolved in the soil domain. Approaches to estimate soil water pressure head drop at the root-soil interface from bulk soil values can be found in Mai et al. (e.g. 2019); Beudez et al. (e.g. 2013). Different approaches for the determination of the sink term for root water uptake are likely to differ most in dry soil. The reference solution to this benchmark is designed to evaluate possible differences between the models in that regard.

##### Reference solution

As no analytical solutions exist for this problem of coupled water flow in the soil-root system, we designed a reference solution with a numerical model that explicitly considers the physical presence of roots in the soil domain, i.e., the soil mesh is highly refined around all roots and water uptake is modelled via boundary conditions at all the root surfaces. Thus, this reference solution does not make any assumptions that are inherent in the definition of the sink terms for root water uptake in the line source-based models. An explicit 3D soil grid is also used in Daly et al. (2017). However here, the soil is additionally coupled to the xylem flow in the root. The root is still modeled as a network of one-dimensional segments (center-line representation). Each segment has a specific radius as specified in the RSML grid file to this benchmark. A three-dimensional representation of the root system is implicitly given by the union of all spheres along the root center-lines. Using this implicit representation a soil grid excluding the root system was generated using the C++ geometry library CGAL (The CGAL Project, 2019). In order to reduce the number of vertices in the mesh, the mesh is locally refined around the root-soil interface. The resulting mesh is available in the Gmsh format (Geuzaine and Remacle, 2009) in the data set. For the evaluation of the radial flux, which is a coupling condition on the soil faces *σ* representing the root-soil interface, we integrate over each face

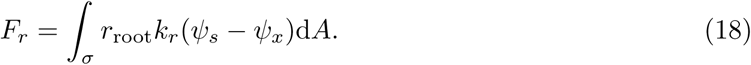

While the soil water pressure head is defined on the face, the corresponding root xylem water pressure head has to be found by a mapping. To this end the integration point is first mapped onto the root surface using its implicit representation. Then the point is mapped onto the corresponding root center-line (a line segment) by finding the closest point on the line segment. There, *ψ*_*x*_ is evaluated. The flux is added as a source term in the corresponding segment in the root. The model is implemented in the open-source porous media simulator DuMu^x^ (Flemisch et al., 2011;.Koch et al., 2019; Koch et al., 2018a). The coupled system is solved with a fully coupled manner, using Newton’s method, and monolithic linear solver (block-preconditioned stabilized bi-conjugate gradient solver) in each Newton iteration. The equations are discretized in time with an implicit Euler method, and in space with a locally mass conservative vertex-centered finite volume method (BOX method (Helmig et al., 1997)). The maximum time step size is Δ*t* = 1200 s. The actual time step size may be sometimes chosen smaller, depending on the convergence speed of the Newton method. Output files are produced in regular intervals every 1200 s starting with the initial solution. The simulation time is 3 d.

Soil water content and root water pressure head in a three-dimensional plot is shown in Fig. 10 for C1.2b. Fig. 11A shows the potential and actual transpiration rates for both scenarios, with constant and age-dependent root hydraulic properties. The curves hardly differ since the water pressure head drop is dominated by the low conductivity of the dry soil. In Fig. 11B, the differences between scenarios are more clearly visible in terms of the minimal and maximal root water pressure head with respect to time.

**Figure 10:**
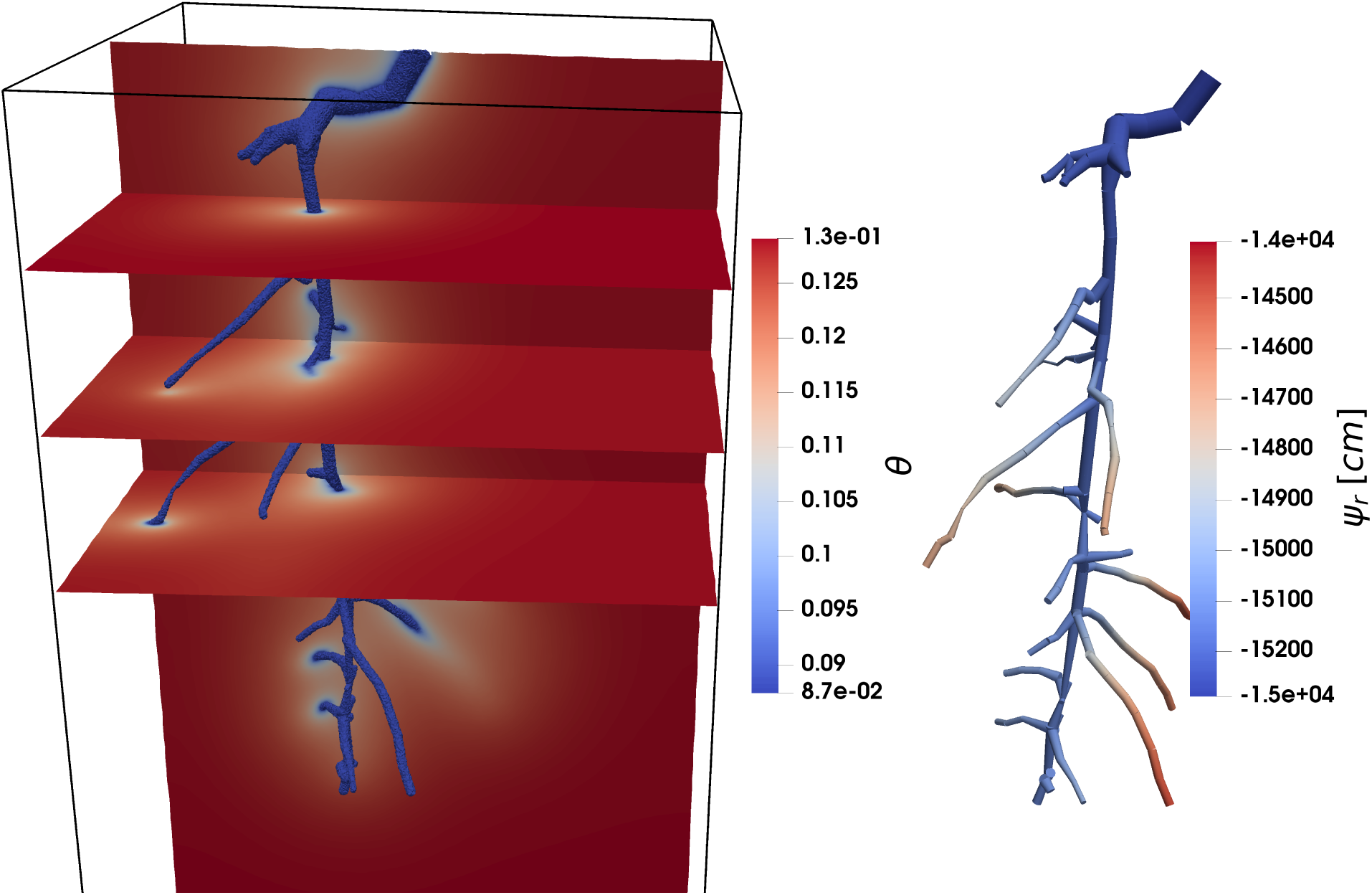
C1.2: Root water uptake by a static root system over time. Soil colours denote volumetric water content, root colours denote root water pressure head.

**Figure 11:**
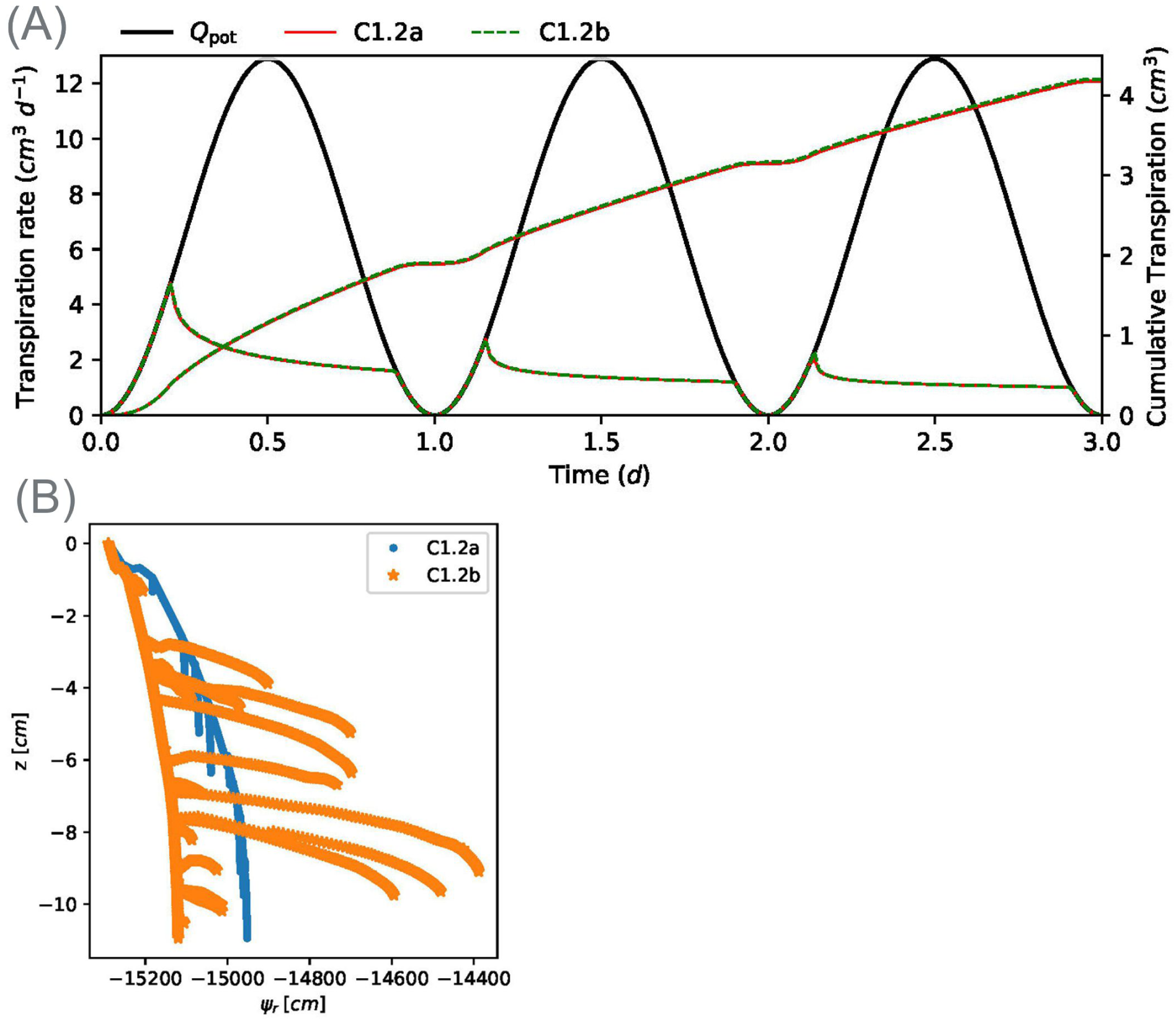
Results of C1.2 for two scenarios, constant and age-dependent root hydraulic properties. A: Actual transpiration of reference solution. B: Root water pressure head distributions inside the root system.

##### Required outputs

To compare the results between the participating models, the desired outputs are

- VTK files (3D) of soil water pressure head and water content on the first, second and third day (*t* = 0.5 d, 1.5 d, 2.5 d). For output written every 1200 s this means the output files with the number 36, 108, and 180.
- VTK files (lines in 3D) of root water pressure head in the first, second and day (*t* = 0.5 d, 1.5 d, 2.5 d)
- CSV file with three data points per time step (each 1200 s starting with *t* = 0): time and actual transpiration rate
- CSV file with three data point per time step: time and minimum and maximum root water pressure head

File names of the VTK files should indicate the simulator name, the state variable, the domain, and the output time in days, e.g. `“DuMux_soil_theta_1d.vtk”.` File names of the CSV files should indicate the simulator name and output time it days, e.g., `“Dumux_1.csv”`.

### 2.2 Coupled benchmark scenarios C2: Root water uptake by a dynamic root system

In this benchmark, we wish to explore differences caused by the approach of root growth modelling. We assess how the differences in root architecture parameters resulting from M1.2 propagate (or not) in the computation of the root water uptake from soil. In this example, we do not consider the effect of soil properties on root growth, but only the differences that arise from the different root systems according to M1.2.

#### 2.2.1 C2.1: Water uptake by a single root

Before looking at the root system, we look at how the implementation of the growth itself affects computed root water uptake for a single root. This scenario is analogous to C1.1, but with a single root growing at an elongation rate of 2 cm/d from 1 to 10 cm length.

##### Required outputs

The required outputs for model intercomparison are

- VTK files of 3D soil water pressure head and water content in soil at a temporal resolution of 1 day up until 60 days (point data)
- VTK files of xylem water pressure head (point data)
- Text files with two lines: time and corresponding actual transpiration

#### 2.2.2 C2.2: Water uptake by a root system

This scenario is the same as C1.2b, but replacing the static root system with a growing root system. The root growth parameters are for each model the results of M1.2; simulations start from a seed and run until a 60 day old root system. The domain size is 25 × 25 × 100 cm, the potential transpiration *Qpot* = 0.5 cm3d−1 is scaled proportional to the root volume divided by the maximal root volume at maturity.

##### Required outputs

- VTK files of 3D soil water pressure head and water content in soil at a temporal resolution of 1 day up until 60 days (point data)
- VTK files of xylem water pressure head (point data)
- Text files with two lines: time and corresponding actual transpiration

File names of the VTK files should indicate the simulator name, the state variable, the domain, and the output time in days, e.g. `“DuMux_soil_theta_1d.vtk”.` File names of the CSV files should indicate the simulator name and output time it days, e.g., `“Dumux_1.csv”.`

### 2.3 Automated comparison within all benchmark problems

Each benchmark folder on the github repository contains a Jupyter Notebook named “Automated comparison”. It provides the analytical solution of the respective benchmark and in addition includes Python code that automatically loads all the outputs of participating models that are provided in the “Numerical results” folder of that benchmark. As soon as new outputs are provided, they are automatically included in the analysis. Currently, different model outputs are already available. We envision more participating models’ outputs to be provided in this way. Future analysis will include graphical and quantitative approaches.

## 3 Conclusions

Functional-structural root architecture models have been compared qualitatively (Dunbabin et al., 2013, e.g.), but until now no quantitative benchmarking existed. In other communities, bench-marking has been done or is ongoing, e.g., AgMIP (Porter et al., 2014) for crop models, CMIP (Eyring et al., 2016) for climate models, subsurface reactive transport models (Steefel et al., 2015). With this paper, we propose a framework for collaborative benchmarking of functional-structural root architecture models that allows quantitative comparison of the outputs of different simulators with reference solutions and with each other. This framework is presented using Jupyter Note-books. Behind every “module” benchmark, there is a working code that explains and implements the reference solution or analysis of reference data. For both, “module” and “coupled” benchmarks, Jupyter Notebooks facilitate the automated comparison of simulator simulation outputs that are stored in specified folders of a public github repository. In this way, new numerical simulators that may be developed in the future may still be added to the automated comparison. All the analysis that is done in the Jupyter Notebooks is freely available so that the comparisons and analysis of uploaded model outputs will be transparent and repeatable. Future efforts will aim at extending the benchmarks from water flow in root and soil systems to further processes such as solute transport, rhizodeposition, etc. We expect that this benchmarking will result in a better understanding of the different models and contribute towards improved models, with which we can simulate various scenarios with greater confidence. It will set standards for future model developments, ensuring that bugs, numerical errors or conceptual misunderstandings do not affect the value of future work. This is a step towards developing those models into the much-needed aid in the design of agricultural management schemes and model-guided crop breeding. These models may also be useful in ecology, e.g. to study species complementarity.

## Acknowledgements

A.S. acknowledges funding by the German Research Foundation (grant number SCHN 1361/3-1). V.C. was supported by the Belgian Fonds National de la Recherche Scientifique (FNRS, grant FC84104). V.S. acknowledges funding by the German Research Foundation (grant number SCHM 997/33-1). This research was institutionally funded by the Helmholtz Association (POF III Program—Research Fields Key Technologies for the Bioeconomy and Terrestrial Environment).

## A Derivation of the analytical solution of water flow inside the root system

The axial water flow rate in the xylem *Q*_*x*_ (cm^3^ day^−1^) is given by

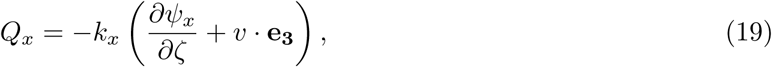

where *k*_*x*_ is the axial conductance (cm^3^ day^−1^), *ψ*_*x*_ is the pressure inside the xylem (cm), *ζ* is the local axial coordinate **e**_**3**_ the unit vector in *z*-direction, and *v* the normalised direction of the xylem.

The radial water flow rate is given by

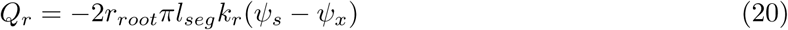

with units (cm^3^ day^−1^), where *r*_*root*_ is the root radius (cm), *l*_*seg*_ is the length of each root segment (cm), *k*_*r*_ is the radial conductivity (day^−1^), and *ψ*_*s*_ is the soil water pressure head of the surrounding soil (cm). The equation is neglecting osmotic potential and is based on Eq. (3.3) of Roose and Fowler (2004). Note that around the root a homogeneous soil water pressure head is assumed, therefore there is actually no hydrostatic equilibrium.

For each segment of length *l*_*seg*_ mass conservation yields

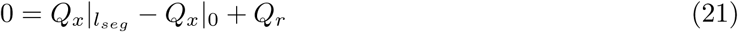

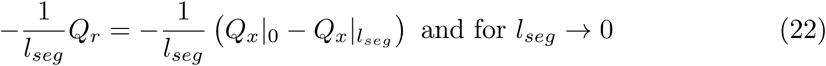

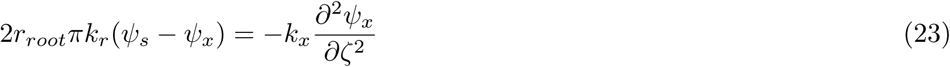

see Eq. (3.4) of Roose and Fowler (2004), where *v*_3_ is the *z*-component of the normalised xylem direction (cm).

Integrating this ordinary differential equation leads to an explicit equation for *ψ*_*x*_(*ζ*)

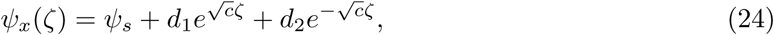

where *c*:= 2*aπk*_*r*_*/k*_*x*_, and *d*_1_, and *d*_2_ are integration constants that are derived from the boundary conditions.

To exemplify, we calculate *d*_1_, and *d*_2_ for a Dirichlet boundary condition at the root collar, and no-flux boundary conditions at the tip. The Dirichlet boundary conditions at the collar of the root system *ψ*_*x*_|_collar_ = *ψ*_0_ is inserted into the analytic solution Eq. (24), and yields

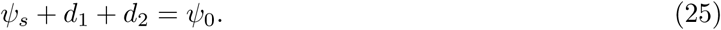

The Neumann boundary condition 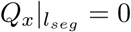 (Eq. 20) leads to

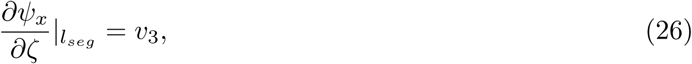

where *l*_*seg*_ is the length of the root segment. Using the derivation of the analytical solution yields

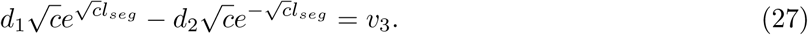

For a straight downward segment *v*_3_ = −1, Eqns (25) and (27) can be summarized as

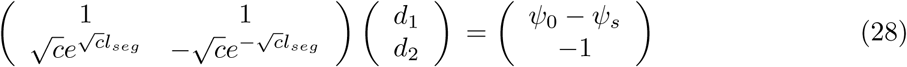

Solving this linear equation for *d*_1_ an *d*_2_ yields

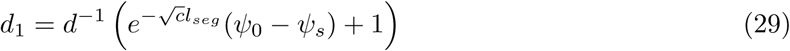

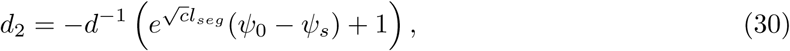

where *d* is the determinant of above matrix

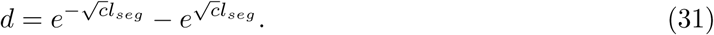

## Notes

https://github.com/RSA-benchmarks/collaborative-comparison

